# Automatic Classification of Prostate Cancer Gleason Scores from Digitized Whole Slide Tissue Biopsies

**DOI:** 10.1101/315648

**Authors:** Hongming Xu, Sunho Park, Tae Hyun Hwang

**Affiliations:** Department of Quantitative Health Sciences, Cleveland Clinic, Cleveland, OH USA

**Keywords:** Prostate cancer, Medical image analysis, Texture descriptor, Feature classification

## Abstract

Histological Gleason grading of tumor patterns is one of the most powerful prognostic predictors in prostate cancer. However, manual analysis and grading performed by pathologists are typically subjective and time-consuming. In this paper, we propose an automatic technique for Gleason grading of prostate cancer from H&E stained whole slide biopsy images using a set of novel completed and statistical local binary pattern (CSLBP) descriptors. First the technique divides the whole slide image into a set of small image tiles, where salient tumor tiles with high nuclei densities are selected for analysis. The CSLBP texture features that encode pixel intensity variations from circularly surrounding neighborhoods are then extracted from salient image tiles to characterize different Gleason patterns. Finally, CSLBP texture features computed from all tiles are integrated and utilized by the multi-class support vector machine (SVM) that assigns patient biopsy with different Gleason score of 6, 7 or ≥8. Experiments have been performed on 312 different patient cases selected from the cancer genome atlas (TCGA) and have achieved more than 79% classification accuracies, which is superior to state-of-the-art textural descriptors for prostate cancer Gleason grading.

## 1 Introduction

Prostate cancer is the second most common cancer in men and the fourth most common tumor type worldwide [1]. Although there have been significant changes in clinical and histologic diagnosis of prostate cancer, the Gleason grading of biopsy tissues remains one of the most powerful prognostic predictors in prostate cancer. Since prostate cancer is a biologically heterogeneous disease with variable molecular alterations, the manual grading by pathologists often suffers from inter- and intra-observer variations. Therefore, the ability to automatically assign Gleason scores from the diagnostic biopsy slide would have significant impacts on clinical decision making, and prediction of patient outcomes.

The Gleason grading system defines five histological patterns from least aggressive (i.e., grade 1) to most aggressive (i.e., grade 5), based on gland structures in the tissue biopsy [4]. Since most tumors typically have two patterns, the original Gleason score is assigned by adding the two most common patterns in a tumor, with scores ranging from 2 to 10 [2]. However, the current application of Gleason grading system has been changed from the original version. Nowadays pathologists mainly assign Gleason scores 6-10 based on the two dominant tumor patterns, since assignment of Gleason scores 2-5 has poor reproducibility and poor correlation with radical prostatectomy grade [2]. Gleason score along with other clinical variables are usually used to create risk stratification for prostate cancer patient management. For example, patients with Gleason score 6 or below is typically considered as low risk. Patients with Gleason score 7 is considered as intermediate risk, and patients with Gleason score 8 or above is considered as high risk. Fig. 1 shows examples of tumors with different Gleason scores. As observed in Fig. 1, the textural appearance of different grade prostate cancers varies from each other due to abnormal changes of gland structures. Based on gland and tumor patterns, there have been a plethora of published studies addressing automatic prostate cancer grading problem. The existing studies can be broadly divided into two main categories: gland-nuclei-based and texture-based approaches.

**Fig. 1:**
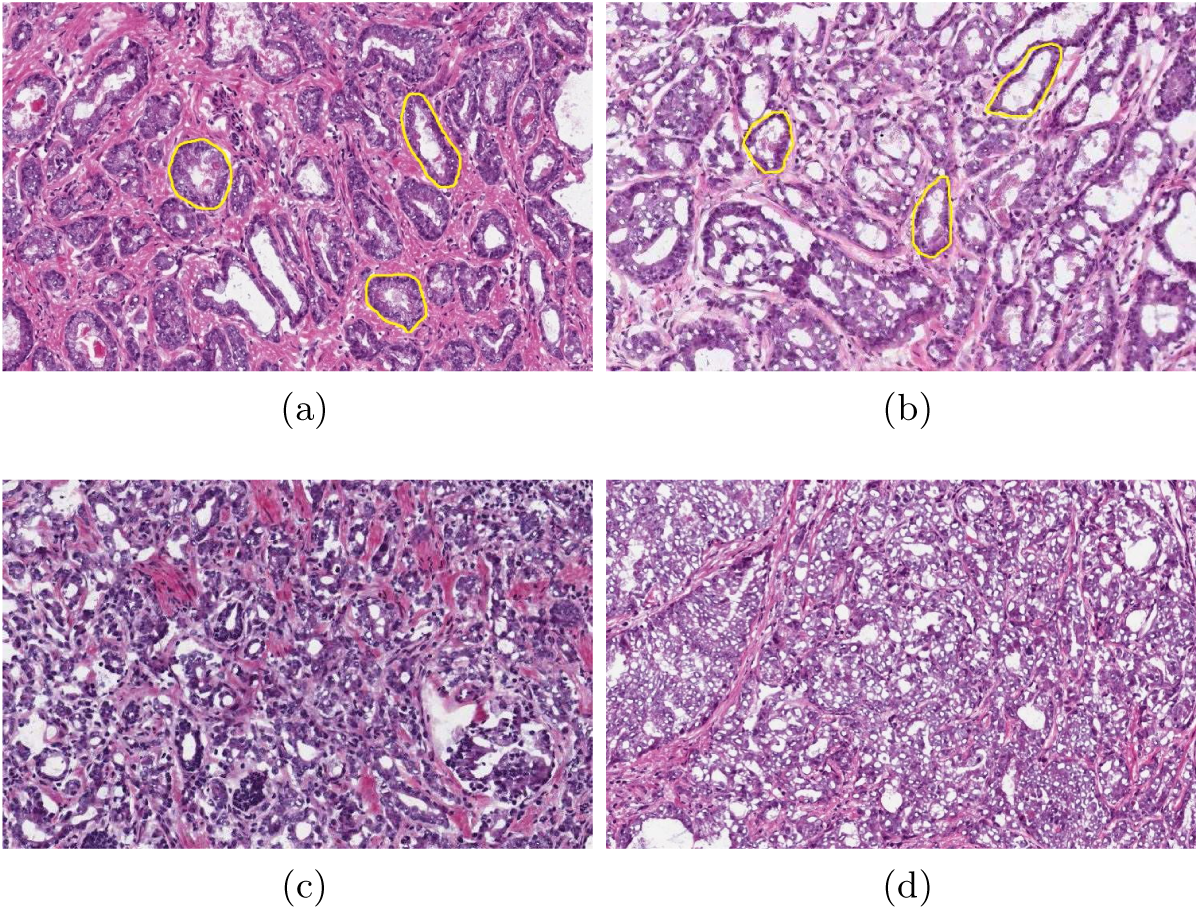
Example of Gleason patterns. (a) Gleason score 6 (low risk). (b) Gleason score 7 (intermediate risk). (c) Gleason score 8 (high risk). (d) Gleason score 9 (high risk). Note that in (a)(b) the superimposed yellow contours highlight a few manually labeled glands.

Gland-nuclei-based techniques try to determine Gleason scores by computing shape and structural information of segmented glands and nuclei in the image. For example, Nguyen et al. [4] proposed a method that first segments glands by incorporating nuclei distribution into a normalized graph cut framework and then assigns Gleason scores based on structural and elliptical features of segmented glands. They reported about 87% accuracies for grade 3 and 4 classifications. Niazi et al. [6] proposed a set of visually meaningful features to differentiate between low (≤ 6) and high (≥ 8) Gleason scores of prostate cancer. These features mainly include the shortest path from epithelial nuclei to the closest luminal and the ratio of epithelial nuclei over the total number of nuclei in the image. They reported a high classification accuracy (above 90%), but the most challenging cases of Gleason score 7 were not included for evaluation. These gland-nuclei-based techniques usually fail to process and identify high Gleason scores, since tumor regions are merged together and there are no clear glands for detection in high Gleason patterns (see Fig. 1(d)).

Texture-based techniques attempt to determine Gleason grade by capturing the textural difference caused by gland changes in different grade of biopsy tissues. The widely-used textural features are computed from gray level co-occurrence matrix, multi-wavelet transform, fractal analysis and texton maps. Huang et al. [3] proposed to classify the prostate biopsy image with different Gleason grades by applying fractal analysis to describe histological variations within the image. Khurd et al. [7] proposed a texture classification technique for Gleason grading, which characterizes the texture in images of different tumor grades by clustering extracted filter responses at each pixel into textons. These texture-based techniques have the advantage that they do not rely on gland or nuclei detections, and hence they are capable of processing high Gleason grade patterns. However, due to high computational complexity of these techniques, they mainly focus on analyzing manually selected regions of interest rather than whole slide biopsy, which is likely to bring sampling bias to prostate cancer grading.

In this paper, we propose an automated technique for Gleason grading of prostate cancer from digitized tissue biopsy slides. There are two main contributions from this work:

– To the best of our knowledge, this is the first study to perform Gleason grading of prostate cancer patients using computerized features from whole histopathology slide image.
– A set of completed and statistical local binary pattern (CSLBP) descriptors are proposed for textural analysis in prostate biopsy images, which are shown to be superior to other traditional textural features for Gleason grading.

## 2 Proposed Method

Fig. 2 depicts an overview of our image processing pipeline. As observed in Fig. 2, the proposed technique mainly consists of three modules. First the whole biopsy slide is preprocessed and divided into a set of non-overlapping image features described by completed and statistical local binary pattern operators are then computed from selected image blocks of the whole biopsy slide. Finally, the selected features are utilized by the classifier, which assigns biopsies to different Gleason scores (i.e., low, intermediate or high risk). In the following subsections, we provide details of our proposed technique.

**Fig. 2:**
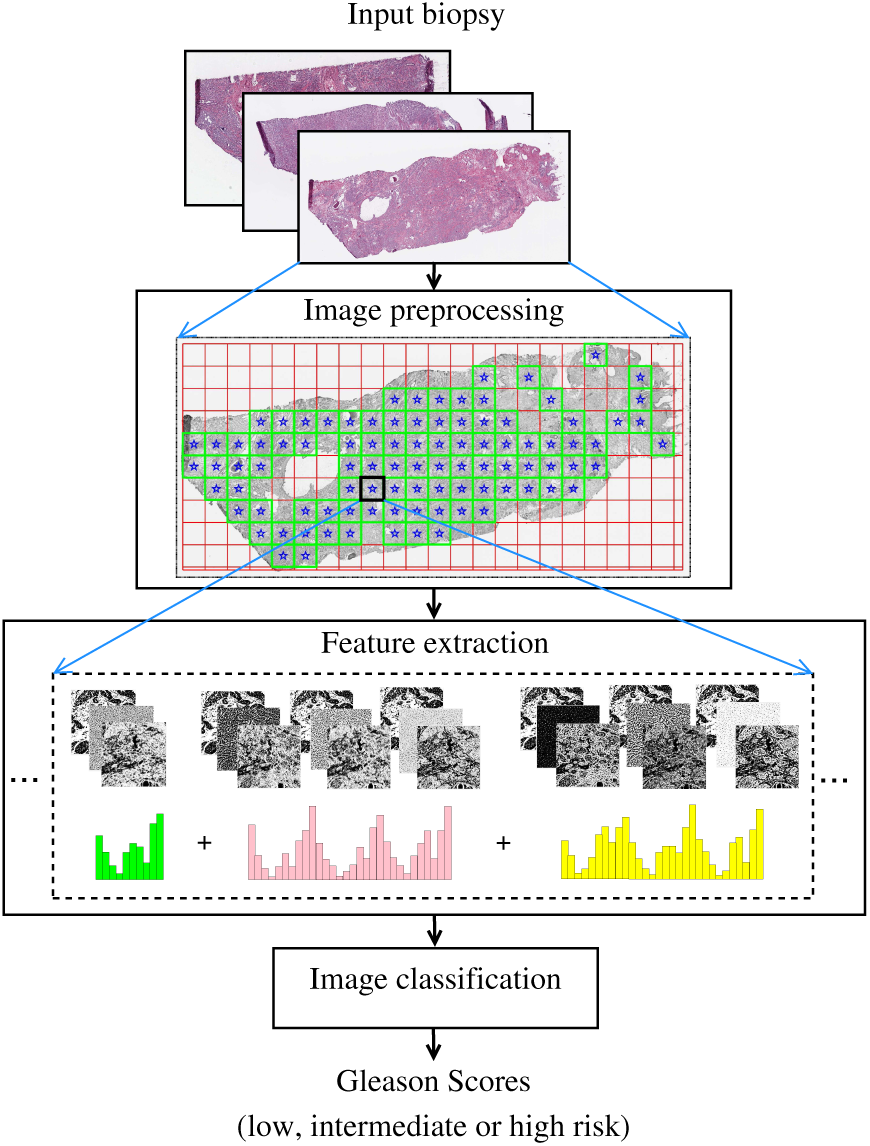
Pipeline of the proposed technique.

### 2.1 Image preprocessing

A whole biopsy slide with a high resolution typically has a large volume size (e.g., about 1GB for 20×), which makes processing it directly with computerized algorithms a challenge [9]. In this module, the biopsy image with 2.5× resolution is selected using the openslide library [11] for efficient processing. The whole biopsy slide is then divided into a series of non-overlapping image blocks that are used by subsequent textural feature analysis. The three steps of this module are as follows.

#### Stain separation

Given an RGB color biopsy image, the color deconvolution method is first applied to separate the image into hematoxylin (H) and eosin (E) channels, respectively [12]. Since most of biological information (e.g., nuclei) is included in H channel, the H channel is then converted into a gray scale image, *H_g_*, for subsequent processing. Fig. 3(a) shows a H&E stained prostate biopsy image, and Fig. 3(b) shows the gray scale image *H_g_*.

**Fig. 3:**
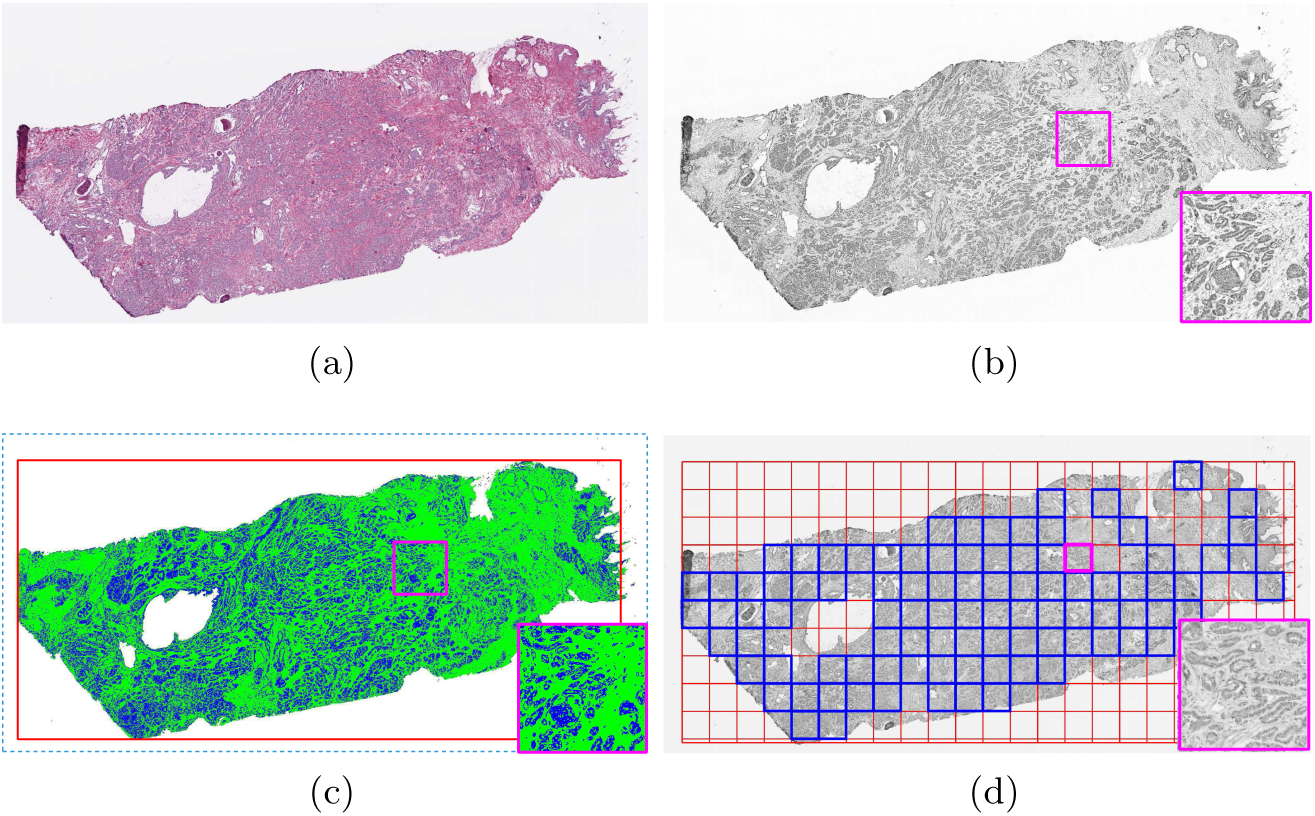
Illustration of image preprocessing. (a) Prostate biopsy slide. (b) Gray scale image *H_g_*. (c) Image segmentations, where white pixels represent image background, green and blue pixels represent stroma and nuclei regions. (d) Image tiling results. Note that in (d) image blocks highlighted by blue squares are selected for subsequent texture analysis.

#### Multi-thresholding

It is observed from Fig. 3(b) that image *H_g_* mainly includes three classes of pixels: background (white), tissue stroma (gray) and nuclei (dark). Based on this observation, the image *H_g_* is segmented into three levels by two thresholds, 0 *< t*_1_ *< t*_2_ *<* 255. The thresholds *t*_1_ and *t*_2_ are automatically determined using Otsu’s method [13], which computes the optimal thresholds by maximizing the inter-class variances. The pixels with intensities below the threshold *t*_1_ are determined as nuclei pixels, while the pixels with intensities below the threshold *t*_2_ are determined as prostate tissue pixels that mainly consist of stroma and nuclei regions. In Fig. 3(c), the white pixels represent segmented image background pixels, whereas green and blue pixels represent segmented stroma and nuclei pixels, respectively.

#### Image tiling

The bounding box [12] of the prostate tissue region (i.e., stroma and nuclei regions) is computed based on image thresholding. In Fig. 3(c) the red rectangle is the determined bounding box for prostate tissue. The image region within the bounding box is then divided into a number of non-overlapping blocks. Let us assume that there are *K* image blocks, and each block has a size of *u* × *v* pixels. The *ith* image block is selected for subsequent feature analysis only if it satisfies the following two inequalities:

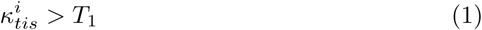

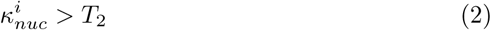

where 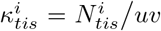 and 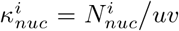. 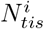 and 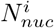 represent the number of prostate tissue and nuclei pixels within the *ith* image block, respectively. *T*_1_, *T*_2_ are two thresholds, which were empirically set as *T*_1_ = 0.9 and 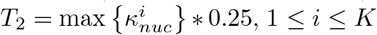, in this work. The above two inequalities are applied such that image blocks containing too many background regions (i.e., *>*10%) or few nuclei pixels are discarded for further feature analysis, as these image blocks tend to include non-tumor regions. The threshold *T*_2_ is adaptively determined as a quarter of the largest nuclei density among all blocks in the image. Note that since nuclei densities vary greatly among different patient biopsies and tumor stages, an adaptive threshold *T*_2_ helps in selecting enough tiles for feature analysis across biopsies with diverse tumor nuclei densities. In Fig. 3(d) red and blue squares highlight divided image blocks, where *u* = *v* = 128. Blue squares in Fig. 3(d) indicate selected image blocks that satisfy the above two inequalities (1)(2). As observed in Fig. 3(d), the blocks with mainly tumor regions are selected for further textural analysis, while the blocks containing too many background or stroma regions have been discarded. Let selected image blocks for further analysis be denoted by *B_i_*, 1 ≤ *i* ≤ *S*, where *S* is the number of selected image blocks.

### 2.2 Feature extraction

After obtaining image blocks from whole slide biopsy, textural analysis is performed to capture different Gleason patterns. Since it would be very time-consuming and also likely to over-grade a tumor at high image magnification, Gleason grading is suggested to perform at a relatively low magnification (e.g., 4x) [2]. Following the recommendation in [2], we map all selected image blocks *B_i_*, 1 ≤ *i* ≤ *S*, from 2.5× to 5.0× magnification for texture analysis in this module. First a set of textural features is computed from every selected image block using proposed completed and statistical local binary pattern (CSLBP) descriptors. The computed textural features from all blocks of the whole biopsy slide are then integrated together to characterize patient tumor severity. The details of this module are described as follows.

#### CSLBP

We first compute CSLBP textural features from every selected image block *B_i_*. The proposed CSLBP is extended from completed local binary pattern (CLBP) [14], and includes one more subcategory termed as statistical local binary pattern (SLBP). In the following, we first briefly describe the CLBP and then detail the SLBP.

##### CLBP

The CLBP is the completed modeling of local binary pattern (LBP) operator [15], which encodes textural information by computing sign and magnitude differences between the central pixel and its surrounding neighbors. Fig. 4 illustrates the computation of CLBP. As observed in Fig. 4, given a central pixel *g_c_* and its *p* circularly (with radius *r*) and evenly spaced neighbors *g_n_*, *n* = 0, 1*, …, p* − 1, the CLBP encoding consists of three components:

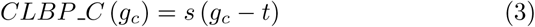

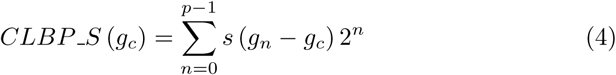

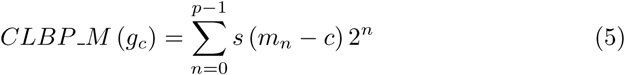

where *CLBP_C* (*g_c_*), *CLBP*_*S* (*g_c_*) and *CLBP*_*M* (*g_c_*) encode the center, sign and magnitude information for pixel *g_c_*, respectively. *s* (·) is the sign function, i.e.,

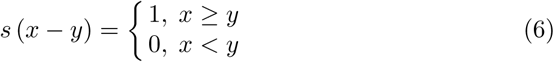

**Fig. 4:**
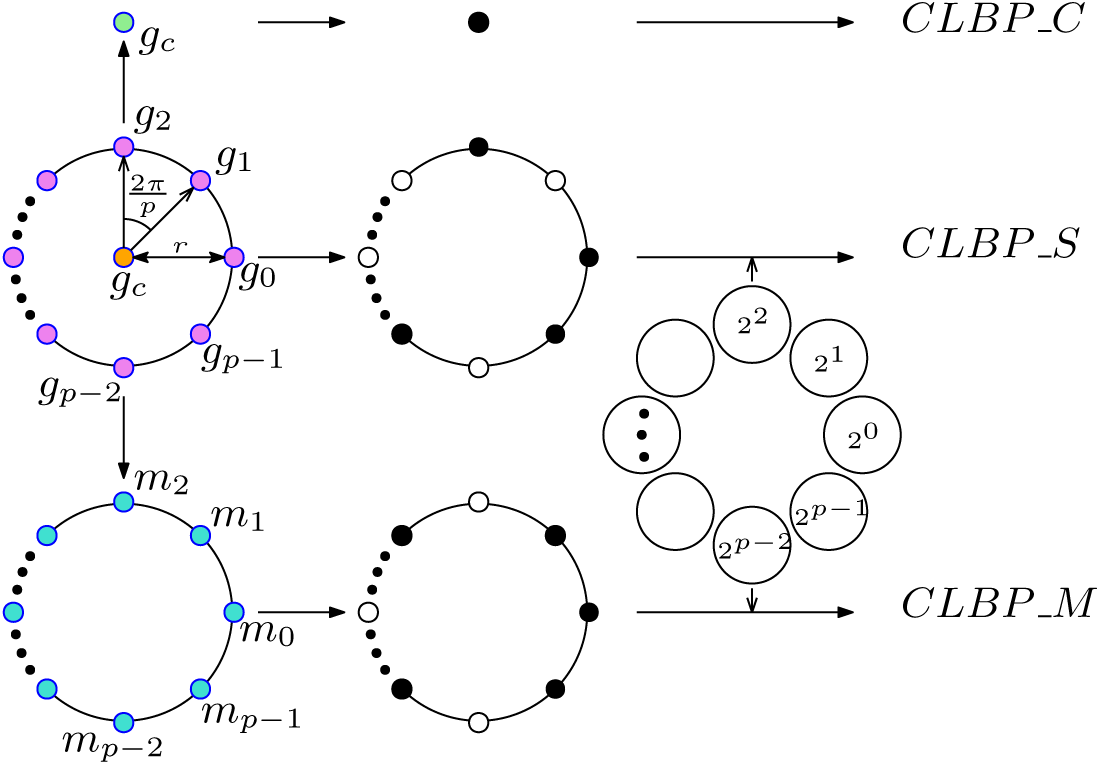
Illustration of completed local binary pattern (CLBP).

The threshold *t* is the average gray intensity of the whole image block (under 5x magnification). *m_n_* is the absolute difference between the center pixel and its neighbor *g_n_*, and c is the mean value of *m_n_* over the whole image block [14].

After computing CLBP S and CLBP M for each pixel in image block *B_i_*, the rotation invariant uniform *riu*2 encoding scheme [15] is applied, which reduces the values of *CLBP_S* and *CLBP_M* from the range 0 ~2^p^ − 1 to 0 ~ p + 1. For more details about *riu*2 encoding scheme, please refer to references [15–17]. In the second row of Fig. 7, it shows three feature maps *CLBP_C*, *CLBP_S* and *CLBP_M*, respectively. To ensure a relatively small feature dimension, a joint 2D histogram *CLBP_M/C* is first built from feature maps *CLBP_C* and *CLBP_M*.The 2D histogram *CLBP_M /C* is then converted to a 1D histogram and concatenated with the histogram of feature map *CLBP_S* to generate a joint histogram, denoted by *CLBP_S_M/C*. In total, (*p* + 2) ∗ 3 histogram features are extracted from CLBP descriptors.

##### SLBP

Although CLBP is capable of capturing microstructure information (e.g., corner and edge) in the image, it is susceptible to image noise and it is difficult to capture macrostructure information due to a small number of neighboring pixels analyzed. To overcome these limitations, we propose SLBP to encode neighboring pixels in a large surrounding region. Formally, given the center pixel *g_c_* and its *p* surrounding neighborhoods *G_n_*, *n* = 0, 1, …, *p* − 1, the *SLBP_C* (*g_c_*), *SLBP_S* (*g_c_*), and *SLBP_M* (*g_c_*) descriptors are computed as follows:

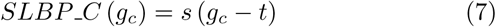

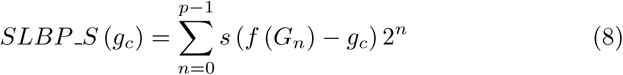

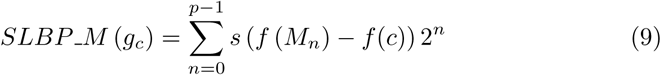

where *SLBP_C* (*g_c_*) encodes the center pixel *g_c_* which is the same as the CLBP shown in Eq. (3). *SLBP_S* (*g_c_*) and *SLBP_M* (*g_c_*) encode the sign and magnitude information of *g_c_* by analyzing its surrounding neighborhoods. *G_n_* represents a set of pixels in the nth neighborhood of *g_c_*, and *M_n_* represents a set of absolute differences between pixels in *G_n_* and *g_c_*. *f* (·) represents a statistical filter function. *f* (*c*) is the threshold and is computed as:

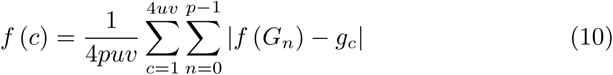

where *u* and *v* are image block size determined during image tiling (see Section 2.1).

Note that different kinds of neighborhood *G_n_* could be explored to analyze surrounding patterns for pixel *g_c_*. In this work, we proposed two surrounding neighborhoods: orientational neighborhood and radial neighborhood. Fig. 5 illustrates SLBP with orientational neighborhood (henceforth referred to as O-SLBP), where the surrounding region of pixel *g_c_* is evenly divided into 8 (i.e., *p* = 8) orientational neighborhoods represented by different color of circles, triangles, squares or diamonds. Fig. 6 illustrates SLBP with radial neighborhood (henceforth referred to as R-SLBP), where the surrounding region of pixel *g_c_* is divided into 4 (i.e., *p* = 4) radial neighborhoods based on the distance between every surrounding pixel and center pixel *g_c_*. As observed in Fig. 5 and Fig. 6, the number of pixels within each neighborhood *G_n_* depends on radius *r* of the surrounding region and the type of neighborhood applied.

**Fig. 5:**
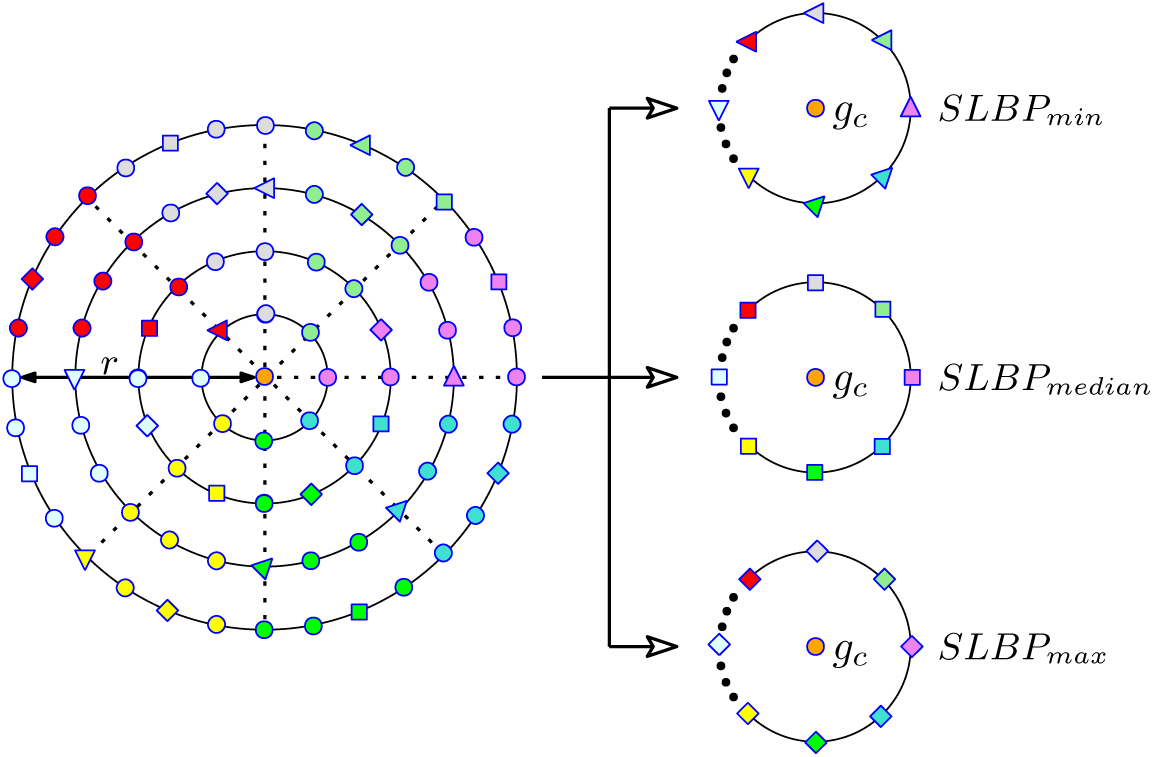
Illustration of statistical local binary pattern with orientational neighborhood (O-SLBP).

**Fig. 6:**
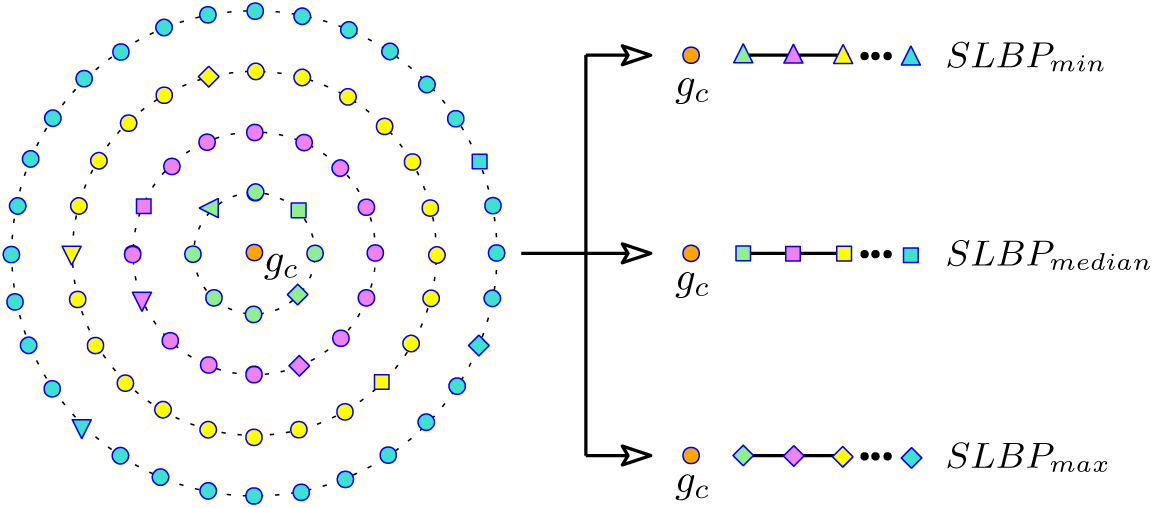
Illustration of statistical local binary pattern (SLBP) with radial neighborhood (R-SLBP).

**Fig. 7:**
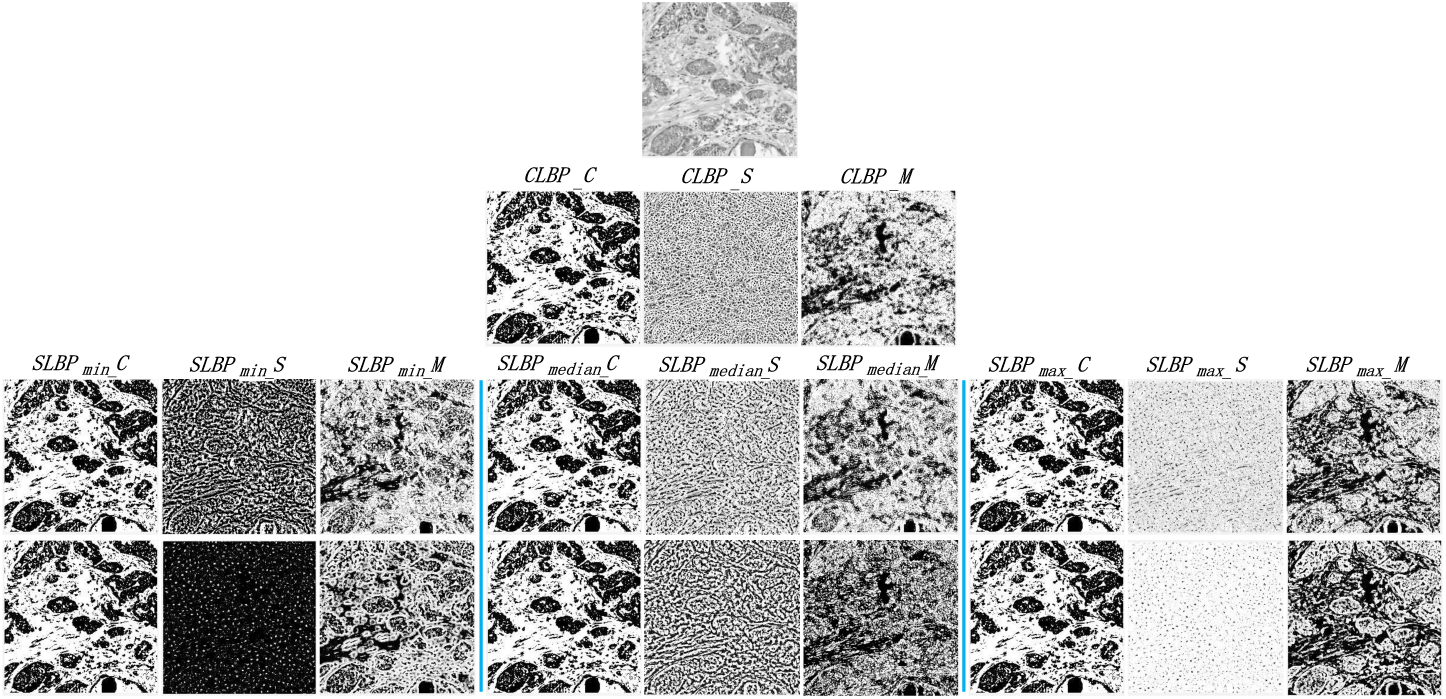
Example of feature maps obtained from CSLBP descriptor. First row: gray input image block; second row: CLBP feature maps; third row: O-SLBP feature maps; fourth row: R-SLBP feature maps.

Similarly, different types of statistical function *f* (·) could be utilized to filter out pixels from the surrounding neighborhood G_n_. In this work, we propose to use three filter functions: min, median and max, to select pixels from neighborhoods *G_n_*, 0 ≤ *n* ≤ *p* − 1. In Figs. 5, 6, the triangles, squares and diamonds represent the minimum, median and maximum pixels in each neighborhood, respectively. As observed in Figs. 5, 6, after selecting the minimum, median and maximum pixels from neighborhoods, three neighboring patterns for the center pixel *g_c_* are generated: *SLBP_min_*, *SLBP_median_* and *SLBP_max_*. These three neighboring patterns are separately encoded with Eqs. (7)(8)(9). For the O-SLBP shown in Fig. 5, the *riu*2 encoding scheme [15] is applied on *SLBP_S* and *SLBP_M*, since neighbors circularly surrounding the center pixel *g_c_* are the same as CLBP. In the third row of Fig. 7, nine feature maps show the corresponding *SLBP_min_, SLBP_median_* and *SLBP_max_* of orientational neighborhoods. After obtaining these feature maps, joint histograms *SLBP_min_*_*S_M/C*, *SLBP_median_*^_^*S_M/C*, and *SLBP_max_^_^S_M/C* are separately computed and concatenated together, which results in (*p* + 2) ∗ 3 ∗ 3 dimensional histogram features together. For the R-SLBP shown in Fig. 6, since statistical neighbors are selected along radial directions and they are invariant regrading image rotations, it is not necessary to further apply the *riu*2 encoding scheme on *SLBP_S* and *SLBP_M*. In the fourth row of Fig. 7, nine feature maps show the corresponding *SLBP_min_*, *SLB*P_*median*_ and *SLBP_max_* of radial neighborhoods. After obtaining these feature maps, joint histograms *SLBP_min_^_^ S*_*M/C*, *SLBP_median_^_^ S*_*M /C, SLBP_max_^_^ S*_*M/C* are separately computed and concatenated together, which results in 2^p^ ∗ 3 ∗ 3 dimensional histogram features together.

#### feature integration

After computing CSLBP texture features from every image block *B_i_*, 1 ≤ *i* ≤ *S*, statistical distribution measures are computed for every CSLBP feature such that feature values across different image blocks of the whole slide biopsy are integrated together. These statistical measures include mean, standard deviation, skewness and kurtosis [5]. Skewness is a measure of the asymmetry of the data around the sample mean. Kurtosis is a measure of whether the data are heavy-tailed or light-tailed relative to a normal distribution. Consequently, after computing 4 statistical measures across *S* image blocks for all CSLBP features, 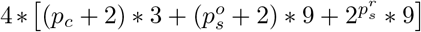 dimensional feature vector per patient biopsy is obtained, where 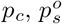 and 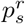 represent the number of surrounding neighbors for CLBP, O-SLBP and R-SLBP, respectively.

### 2.3 Image classification

With computed textural features from whole slide biopsy, the patient tumor is now classified into different categories of Gleason patterns using machine learning classifiers. In this work, principal component analysis (PCA) is first applied to reduce feature dimensions [5]. The PCA is applied for two main reasons. (1) high dimensional features are reduced into low dimensional space in order to prevent over-fitting to the training dataset. (2) PCA helps reduce feature noise and remove redundant features without contributing much to the discrimination power. Feature standardization is then performed on each feature to make its values have zero mean and unit variance, so that each feature contributes equally to classification. Finally, the “one-against-one” multi-class support vector machine (SVM) method [12] is used to perform prostate cancer patient classification. Let *k* denote the number of classes for classification. In the multi-class SVM method, we construct *k* (*k* − 1)/2 SVM classifiers, and each classifier is trained on data from two classes during the training phase. After obtaining *k* (*k* − 1)/2 SVM classifiers, the test patient image will be labeled by all of them in the testing phase. The test patient is predicted to be in the class that is labeled by most of the binary SVM classifiers.

## 3 Experiments and evaluations

In this section, we evaluate the performance of the proposed technique. First we illustrate the datasets used in this work. We then explain comparisons and evaluation metrics. Finally, the quantitative evaluations and comparisons for Gleason grading of prostate cancer patients are provided.

### 3.1 Image dataset

All whole slide prostate biopsy images were obtained from the cancer genome atlas (TCGA) [1], which were stained with hematoxylin and eosin (H&E). TCGA prostate data was derived from multiple institutions over many years, and includes a cohort of 496 prostate adenocarcinoma patients in total. However,TCGA biopsy slides are scanned from frozen tissue sections, so some images have relatively poor quality due to tissue distortions. These slides are not suitable for pathological image analysis. Based on image preprocessing and visual examinations, 312 acceptable patient biopsy slides are used in this study. Note that although one patient may have two or more satisfied biopsy slides, only one patient slide is selected such that no same patient is used in both training and testing during evaluation. The ground truths of Gleason scores are also provided by TCGA on patient records. Among 312 patient slides, 32 patients are diagnosed as low risk with Gleason score 6, 141 patients are diagnosed as intermediate risk with Gleason score 7 and 139 patients are diagnosed as high risk with Gleason score ≥ 8. Since the number of low risk patients (i.e., with Gleason score 6) is far less than that of intermediate or high risk patients, we augment the number of biopsy images for low risk patients. Specifically, after computing the bounding box from prostate tissue pixels for every patient biopsy (see red rectangle shown in Fig. 3(c)), the bounding box is shifted towards bottom-right direction by 30,60 and 90 pixels, respectively. The image tiling in Section 2.1 is then separately performed based on shifted bounding boxes. Thus 1 patient biopsy is augmented to 4 biopsy images, and low risk patient biopsies are increased to 128 images after augmentation. Note that the bounding box shifting has altered selected tumor tiles from the same patient biopsy.

### 3.2 Parameter settings and comparisons

There are two key parameters for CSLBP texture descriptors: radius *r* and number of neighbors *p*. Table 1 illustrates parameter settings for (*r, p*) values of CLBP and SLBP descriptors. Note that large (*r, p*) values would result in high computational complexity and feature dimensions, while (*r, p*) values that are too small may fail to capture macro-structure information in the image. These (*r, p*) values are determined by balancing feature representations and computational complexities. As observed in Table 1, 288 CSLBP textural features are computed from every selected image block of the whole slide image. After computing 4 statistical measures across all selected image blocks (see Section 2.2), one patient biopsy slide corresponds to 1152 feature dimensions. These 1152 feature dimensions are heuristically reduced to 62 dimensions by PCA, where about 85% data variance is retained for evaluations.

**Table 1:** Parameter settings for texture analysis

| Descriptors | Radius $r$ | Neighbors $p$ | Feature dimensions |
| --- | --- | --- | --- |
| CLBP | 2 | 16 | 54 |
| O-SLBP | 4 | 8 | 90 |
| R-SLBP | 4 | 4 | 144 |

To verify the efficacy of proposed CSLBP descriptor for prostate cancer Gleason grading, the proposed technique is compared to several widely-used texture descriptors for medical image analysis. First, we compared our technique with a fractal analysis based technique for prostate cancer Gleason grading [3], henceforth referred to as FA technique. In the FA technique, four groups of grid size including {2, 4, 8}, {8, 16, 32}, {32, 64, 128} and {2, 4, 8, 16, 32, 64, 128} are used for fractal analysis by the differential box-counting method. 4 fractal dimension texture features are derived from image intensity difference and image entropy, respectively, which results in 8 texture features for each selected image block. The mean, standard deviation, skewness and kurtosis of each feature are then computed across all image blocks of the whole slide image, therefore each whole slide biopsy corresponds to 32 dimensional features. These 32 features are used by SVM classifier for prostate cancer grading. Besides the FA technique [3], we adaptively implemented another textural analysis based technique, henceforth referred to as HHG technique, which computes histogram, Haralick and Gabor filter related texture features for prostate cancer grading. These textural features have been used for recognition of different tissue types from colorectal cancer histology images [10] and prostate cancer Gleason grading from magnetic resonance images (MRI) [8]. In our implementation, we first compute 6 first order histogram features, 60 second order Haralick features and 12 Gabor filter related features from every selected image block. The mean, standard deviation, skewness and kurtosis of each feature are then computed across all image blocks of the whole slide image, which results in 312 dimensional features for each patient biopsy. For comparison, these 312 dimensional features are reduced to 62 dimensions by the PCA transform, which are finally used by the SVM for prostate cancer Gleason grading. Note that for all techniques the same default parameter settings of SVM classifier in Matlab toolbox [12] are used for comparison.

### 3.3 Evaluation metrics

In this study, prostate cancer patients are tested with 3 risk groups: low risk (Gleason score 6), intermediate risk (Gleason score 7) and high risk (Gleason score 8 and above) [2]. For evaluation, if the automatic grading is the same as the expert class label, it is considered as a correct classification. The performance of the proposed technique is evaluated by the classification accuracy *ACC*, which is defined as:

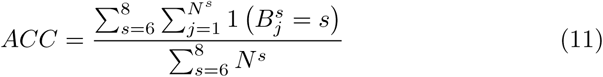

where *N ^s^* is the number of samples belonging to ground truth class *s*, 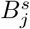 represents the automatically classified result for the *jth* sample in the class *s* and 1 (·) is the indicator function, i.e.,

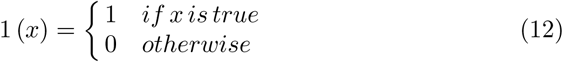

We also compute the area under the receiver operating characteristic (ROC) curve to evaluate the performance of our proposed technique. Since we are dealing with a three-class classification problem, three binary classifications are created for generating ROC curves. Specifically, each time we consider one class as positive samples and the remaining two classes as negative samples to generate a ROC curve. This process is repeated three times and thus three ROC curves are obtained. Let *AU C_k_* denote the area under the *kth* ROC curve, where 1 ≤ *k* ≤ 3. The mean *AU C* for the three ROC curves is computed as:

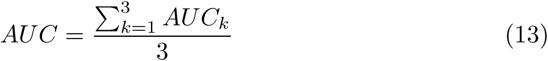

By using the above two evaluation criteria, we perform the 5-fold cross validation and repeat it 50 times for evaluation. The average values of different evaluation criteria are used as the final results.

### 3.4 Classification results

Table 2 lists *ACC* and *AU C* values of prostate cancer Gleason grading by using the FA [3], HHG [10, 8] and proposed CSLBP techniques, respectively. Note that the SVM classifier used by each technique has been tested with polynomial and Gaussian kernels in our evaluations. As observed in Table 2, the FA and HHG techniques provide much poorer performances than the proposed CSLBP technique. These two techniques achieve relatively better performances when the polynomial kernel is applied for SVM classification. Specifically, the FA technique achieves 0.6115 and 0.7833 *ACC* and *AU C* values, while the HHG technique achieves 0.6864 and 0.8544, respectively. In comparison, the proposed CSLBP technique achieves much better performances than the two existing techniques, achieving 0.7908 and 0.9334 *ACC*, *AU C* values when the Gaussian kernel is applied. Additionally, note that the proposed CSLBP technique provides very similar performances when either polynomial or Gaussian kernel is applied, which indicates the stability of our proposed CSLBP descriptors in prostate cancer Gleason grading. The good performance of our proposed technique is mainly because CSLBP descriptors can capture both micro- and macro-texture information by analyzing a large surrounding neighborhood of pixels in the image.

**Table 2:** Comparison of different techniques

| Techniques | | $ACC$ | $AUC$ |
| --- | --- | --- | --- |
| FA [3] | Polynomial | 0.6115 | 0.7833 |
|  | Gaussian | 0.5782 | 0.7733 |
| HHG [10, 8] | Polynomial | 0.6864 | 0.8544 |
|  | Gaussian | 0.6285 | 0.8178 |
| Proposed CSLBP | Polynomial | 0.7850 | 0.9221 |
|  | Gaussian | 0.7908 | 0.9334 |

Fig. 8 shows three ROC curves obtained by different techniques with polynomial and Gaussian kernels for classification. In Fig. 8(a) the low risk group is considered as the positive class, while the intermediate and high risk groups are considered as the negative class. Similarly, in Figs. 8(b)(c) the intermediate and high risk groups are separately considered as positive class, while the remaining two risk groups are considered as negative class. As observed in Fig. 8 the proposed CSLBP technique using either polynomial or Gaussian kernels have provided remarkably better performances than the two existing techniques. In particular, in Fig. 8(a) the areas under the ROC curve for three techniques are much larger than those correspondingly shown in Figs. 8(b)(c). This is mainly because 128 low risk group samples are obtained by augmenting 32 patient digitized biopsies, which results in a relatively good performance for low risk group detection during cross validation. Fig. 9 shows confusion matrices of different techniques for three class predictions. Note that in Fig. 9 the intermediate risk group has been abbreviated as ITM for illustration. Overall, it can be concluded from Fig. 8 and Fig. 9 that the proposed CSLBP descriptors can more accurately distinguish different prostate Gleason patterns than traditional features such as Haralick textures, Gabor filters or fractal analysis as used in [3, 10, 8].

**Fig. 8:**
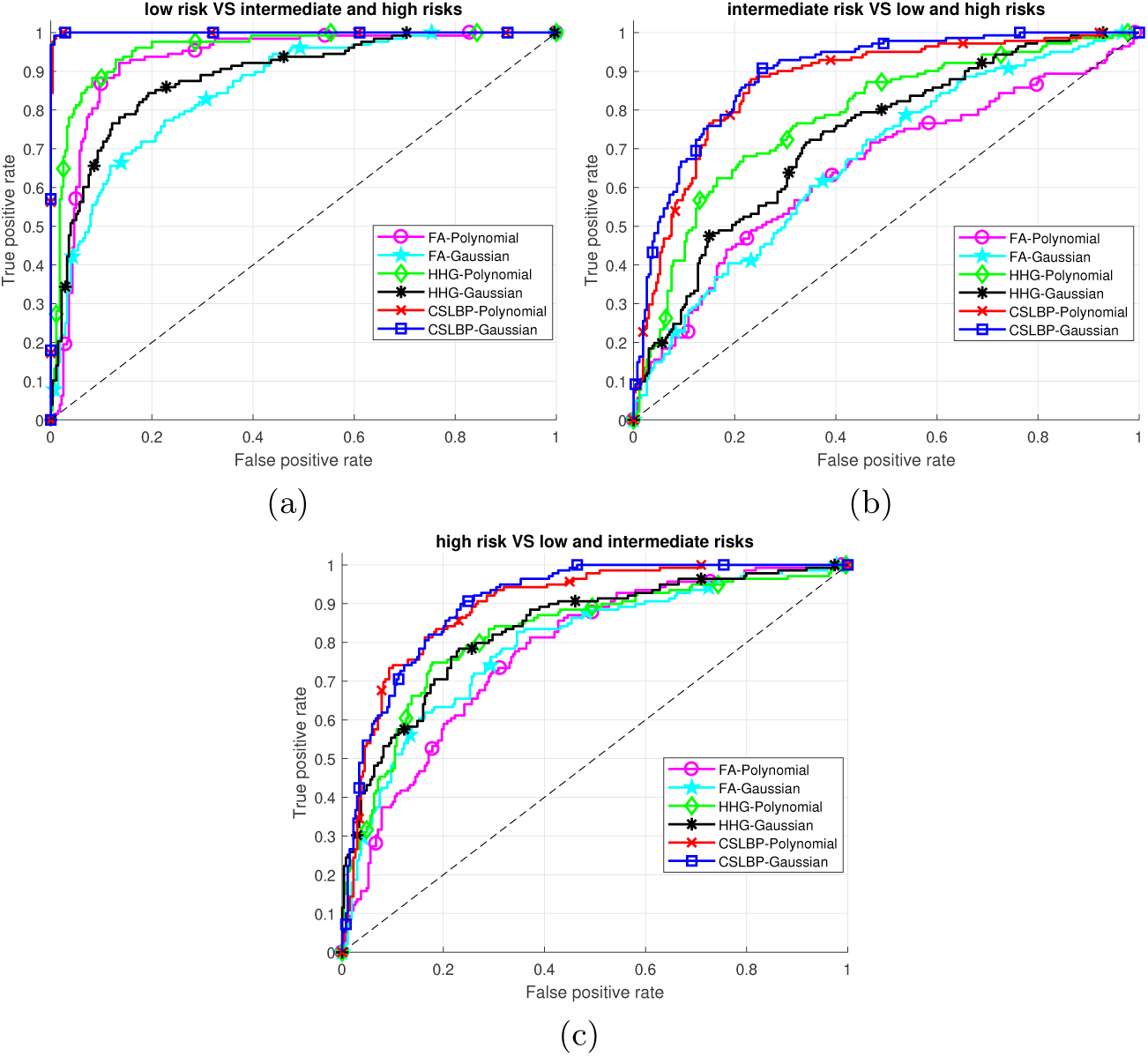
ROC curves for different techniques. (a) Low risk VS intermediate and high risk groups. (b) Intermediate risk VS low and high risk groups. (c) High risk VS low and intermediate risk groups.

**Fig. 9:**
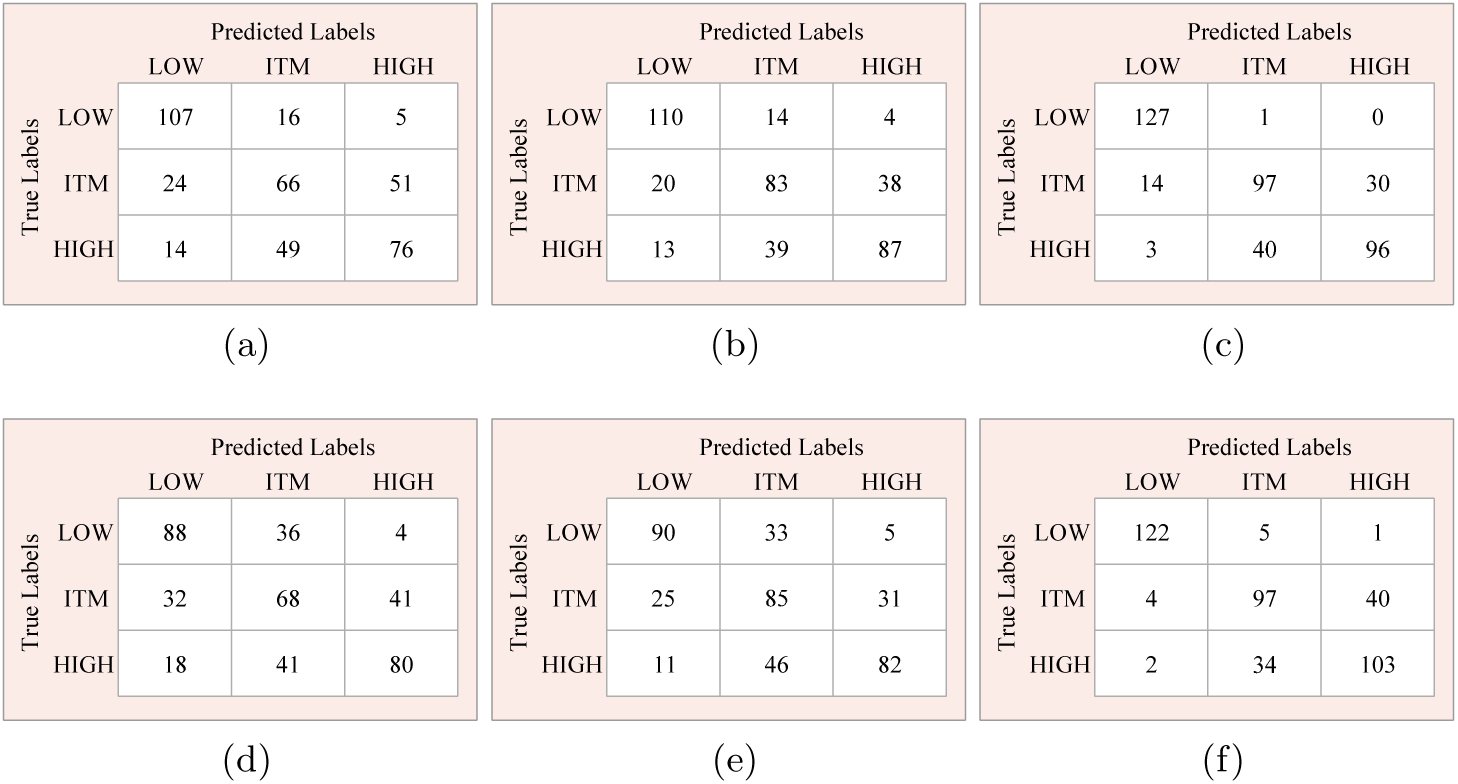
Comparison of confusion matrices for prostate cancer grading. (a) FA with polynomial kernel. (b) HHG with polynomial kernel. (c) Proposed CSLBP with polynomial kernel. (d) FA with gaussian kernel. (e) HHG with gaussian kernel. (f) Proposed CSLBP with gaussian kernel.

## 4 Conclusions

In this paper, we present an automatic technique for prostate cancer Gleason grading from H&E stained whole slide biopsy images. The technique first divides the whole slide image into a series of image blocks for feature analysis. A set of textural features based on our proposed CSLBP descriptors are then computed from every selected image block and statistically integrated together across all image blocks to form a feature representation for a given patient tumor biopsy. Finally the multi-class SVM classifier is applied to predict patient biopsy into different risk groups corresponding to different Gleason scores. The proposed technique has been evaluated on more than 300 patients from the publicly available TCGA dataset and achieves more than 79% classification accuracy for three risk groups stratification. This indicates the efficacy of the proposed CSLBP descriptors for prostate cancer Gleason grading. In future, we will utilize our proposed CSLBP descriptors to explore other cancer types for clinical outcome prediction.

